# *Salmonella*-driven Intestinal Edema in Mice is Characterized by Tensed Fibronectin Fibers

**DOI:** 10.1101/2023.07.26.550515

**Authors:** Ronja Rappold, Konstantinos Kalogeropoulos, Ulrich auf dem Keller, Viola Vogel, Emma Slack

**Affiliations:** Institute of Translational Medicine, ETH Zurich, Zurich, Switzerland; Department of Biotechnology and Biomedicine, Technical University of Denmark, Kgs. Lyngby, Denmark; Institute of Food, Nutrition and Health, ETH Zurich, Zurich, Switzerland; Botnar Research Center for Child Health, Basel, Switzerland

## Abstract

Intestinal edema is a common manifestation of numerous gastrointestinal diseases and is characterized by the accumulation of fluid in the interstitial space of the intestinal wall. Technical advances in laser capture microdissection and low-biomass proteomics now allow us to specifically characterize the intestinal edema proteome. Our data identifies a high abundance of antimicrobial factors, but also highlights major contributions from the blood clotting system, extracellular matrix (ECM) and protease – protease inhibitor networks. The ECM is a complex fibrillar network of macromolecules that provides structural and mechanical support to the intestinal tissue. One abundant component of the ECM observed in the *Salmonella*-driven intestinal edema is the mechanosensitive glycoprotein fibronectin. Using mechanosensitive staining of fibronectin reveals a solely tensed molecular conformation of fibronectin fibers present in the edema, in contrast to other cecal tissue areas, despite the high abundance of proteases able to cleave fibronectin. Co-staining for fibrin(ogen) indicates the formation of a provisional matrix, similar to what is observed in response to skin injury. The absence of low tensional fibronectin fibers and the additional finding of a high number of protease inhibitors in the edema proteome could indicate a critical role of stretched fibronectin fibers in maintaining tissue integrity in the severely inflamed cecum. Understanding these processes may provide valuable functional diagnostic markers of intestinal disease progression.

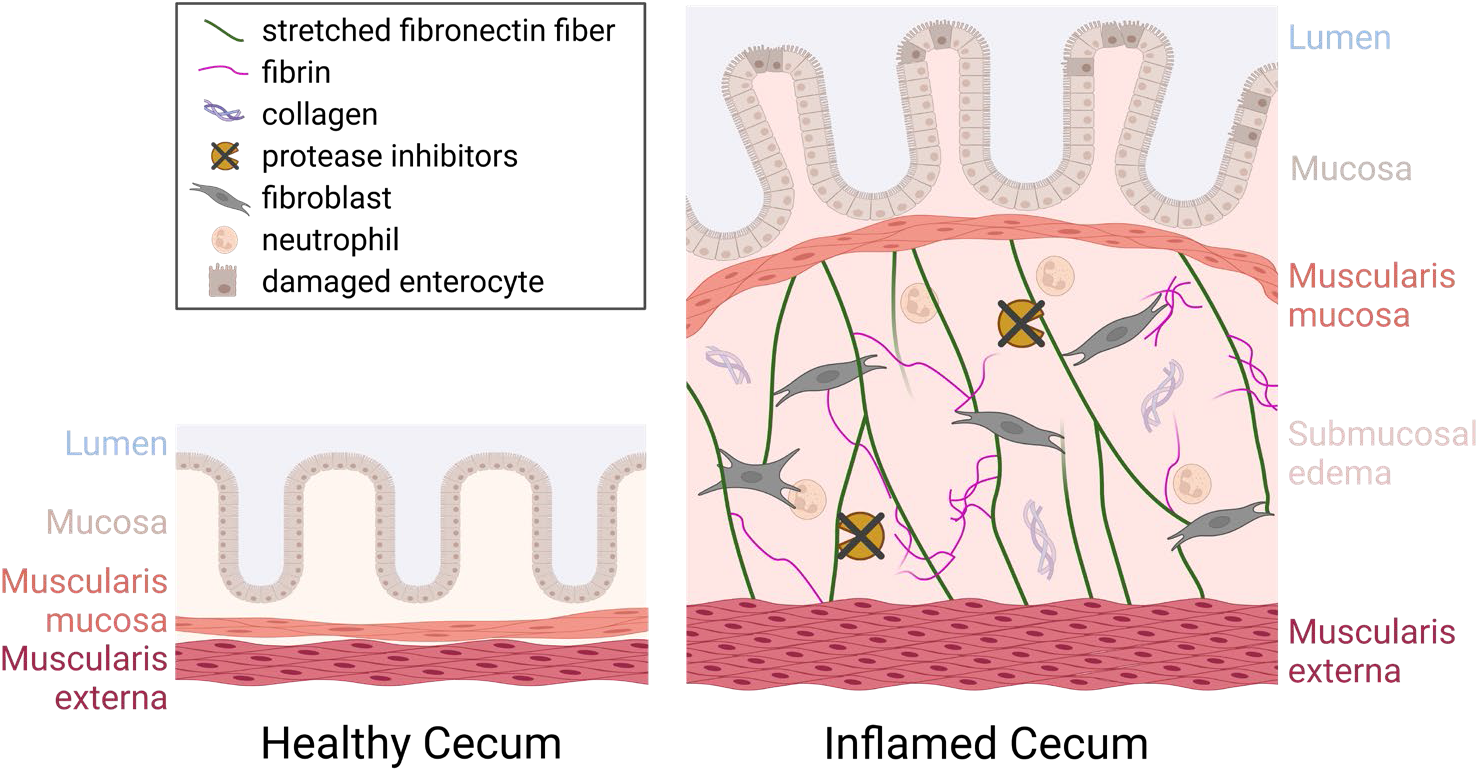

## Introduction

The function of the gastrointestinal tract is to digest and absorb nutrients from food into the mammalian body, while maintaining the physical as well as immunological barrier against harmful substances and pathogens from outside. During an invasive enterobacterial infection, local and systemic inflammatory reactions act to limit the size of the invaded pathogen population and to prevent its spread to systemic sites. Intestinal edema is caused by the build-up of excess fluid in the interstitial spaces of the intestinal wall. Edema formation together with immune cell infiltration and ulceration of the epithelial layer are common pathological observations in intestinal diseases (Erben et al., 2014). Abnormal fluid accumulation in the interstitium of the intestinal wall tissue leads to swelling and physical separation of the two smooth muscle layers present in the cecum. Intestinal edema formation is typically thought of as detrimental to tissue homeostasis due to the impairment of the organ’s mechanical and physiological functions. The diffusion distance of oxygen and other nutrients is dramatically increased, while the transport of toxic byproducts is impeded (Moore-Olufemi et al., 2005). Furthermore, it was reported that edema can lead to a decrease in smooth muscle cell contraction in the intestine, limiting bowel movement in the already challenged inflamed organ (Moore-Olufemi et al., 2009; Uray et al., 2006). Conversely, edema represents a useful response to intestinal infection as tissue swelling reduces the gut luminal volume, limiting the colonizable space, and therefore the total population size, for pathogens such as *Salmonella* Typhimurium. Non-typhoidal *Salmonella* infections are still one of the most common causes of diarrhoeal diseases and represent a major public health concern with several tens of million cases of gastroenteritis globally each year (Gong et al., 2022; Majowicz et al., 2010; World Health Organization, 2018). One of the most important subspecies is *Salmonella enterica* subspecies *enterica* serova Typhimurium (*S.* Tm), causing gastroenteritis in humans and animals. *S.* Tm infection mechanisms have been widely studied (Hausmann and Hardt, 2019; Wotzka et al., 2017) using germ-free, or antibiotics pre-treated, specific-pathogen free mice (Barthel et al., 2003; Stecher et al., 2005). After oral infection, *S.* Tm passes through the gastrointestinal tract and reaches the large intestine, its major site of replication in the mouse intestine (Furter et al., 2019). Tissue invasion by the bacteria induces a multitude of host defense mechanisms: secretion of antimicrobial peptides, recruitment of neutrophils and other inflammatory cell types, expulsion of infected epithelial cells as well as production of pathogen-specific IgA to clear bacteria from the lumen (Hausmann et al., 2020; Maier et al., 2014; Moor et al., 2017; Muniz et al., 2012). These host reactions limit systemic spread of *Salmonella* (Hausmann and Hardt, 2019), but also cause collateral damage to the microbiota and damage to intestinal tissues (Fattinger et al., 2021; Stecher et al., 2007).

The extracellular matrix (ECM), the complex network of proteins and carbohydrates providing structural integrity to cells and tissues, is now considered to play an important role in controlling interstitial volume (Stewart, 2020). However, it is still unclear how exactly the ECM contributes or counteracts potential tissue swelling. The main ECM components include collagens, glycoproteins such as fibronectin, and carbohydrates such as glycosaminoglycans, and various ECM-associated growth factors (Pompili et al., 2021). In particular, fibronectin’s important role in tissue growth and wound healing has been extensively studied (Hynes, 2009; Lenselink, 2015; Patten and Wang, 2020). Fibronectin knock-out mice do not survive past early stages of development and the embryos start to deteriorate after embryonic day 8 (George et al., 1993). This lethality demonstrates the vital role that fibronectin plays in the overall development and survival of animals and limits animal studies to active measurement and observation. Fibronectin fibers as well as a variety of other ECM fibers can be mechanically stretched and partially unfolded, entailing functional consequences such as altered growth factor or cytokine binding, altered enzymatic fiber cleavage or accessibility to binding sites (Vogel, 2018). Fibronectin fiber stretching can also activate cryptic bindings sites, which are not accessible in the equilibrated fibronectin molecule and are important for, but not limited to, fibronectin matrix assembly (Hynes, 2009; Vogel, 2018). Additionally, collagen I fibrillogenesis and interleukin-7 binding have been reported to be dependent on the tensile state of fibronectin fibers (Franyuti et al., 2017; Kubow et al., 2015), which in turn also was shown to affect cellular signaling and behavior *in vitro* (Li et al., 2013; Smith et al., 2007). Nevertheless, if and how fibronectin fibers are involved in the build-up or resolving phase of intestinal edema remains to be investigated.

Studying edema composition is technically challenging as there are few cells, rendering spatial transcriptomics uninformative. There are no known specific reporter or cell-type-specific knock-out systems for interventional mechanistic studies, necessarily limiting us to detailed observation. Additionally, the contribution of these fluid-filled areas to the total tissue proteome is minimal, such that standard proteomics approaches fail to represent the edema content. Recent advances in microscopy, laser capture microdissection and proteomics on very-low biomass samples now make this possible. In this *in vivo* study, we explore the biochemical changes occurring during *Salmonella-*driven edema formation and investigate how fibronectin fiber stretching changes during the tissue remodeling process. We are able to address this question for the first time, using a fibronectin fiber tension probe that binds only to low tension, but not to high tension fibronectin fibers. Unbiased discovery proteomics of dissected edema tissue reveals the protein composition of intestinal edema, and uncovers the major role of the blood clotting system and extracellular matrix within these sites. With this distinct characterization of the edema-specific proteome, we set the foundation for future work on understanding the impact of edema components on cecal inflammation as well as intestinal tissue healing in case of disease survival.

## Results

The streptomycin mouse model for non-typhoidal *Salmonella* infection is a widely used and highly reproducible model of gastrointestinal inflammation, inducing clinical and histological features of typhlocolitis. The transient clearance of gut microbiota by orally administered antibiotics opens a metabolic niche for *Salmonella* in the upper large intestine, allowing *S.* Tm to colonize at densities exceeding 10^9^ colony forming units (CFU) per gram of gut content. The pathological tissue changes of the disease include neutrophil infiltration, submucosal edema, goblet cell depletion and epithelial disruption. Of these phenomena, the induction and control of submucosal edema is the least well-understood. We therefore first characterized the histopathological and mechanobiological changes in *S.* Tm-infected cecal tissue to gain insight into potential mechanoregulatory pathways involved in this process.

25 wildtype specific-pathogen-free C57BL/6 mice were fed a high dose of streptomycin the day before infection. Five of the streptomycin-pretreated animals were mock infected and 20 animals were fed 5×10^7^ CFU of *S.* Tm, mimicking foodborne infection. The widely used inflammation marker, fecal lipocalin-2 (LCN2), was used to determine the severity of the intestinal inflammation directly from feces samples. Fecal LCN2 levels measured over the disease time course confirmed similar disease kinetics in comparison to previously reported infections using oral gavage (Barthel et al., 2003; Maier et al., 2013; Nguyen et al., 2020). The LCN2 values rapidly increased between 6 h post infection (p.i.), and 24 h p.i., up to values of 10^3^ ng/g of feces. Over the next 2 days the LCN2 levels were further raised to 10^4^ ng/g of feces at 72 h p.i., indicating severe gastrointestinal inflammation (Fig.1C).

**Figure 1:**
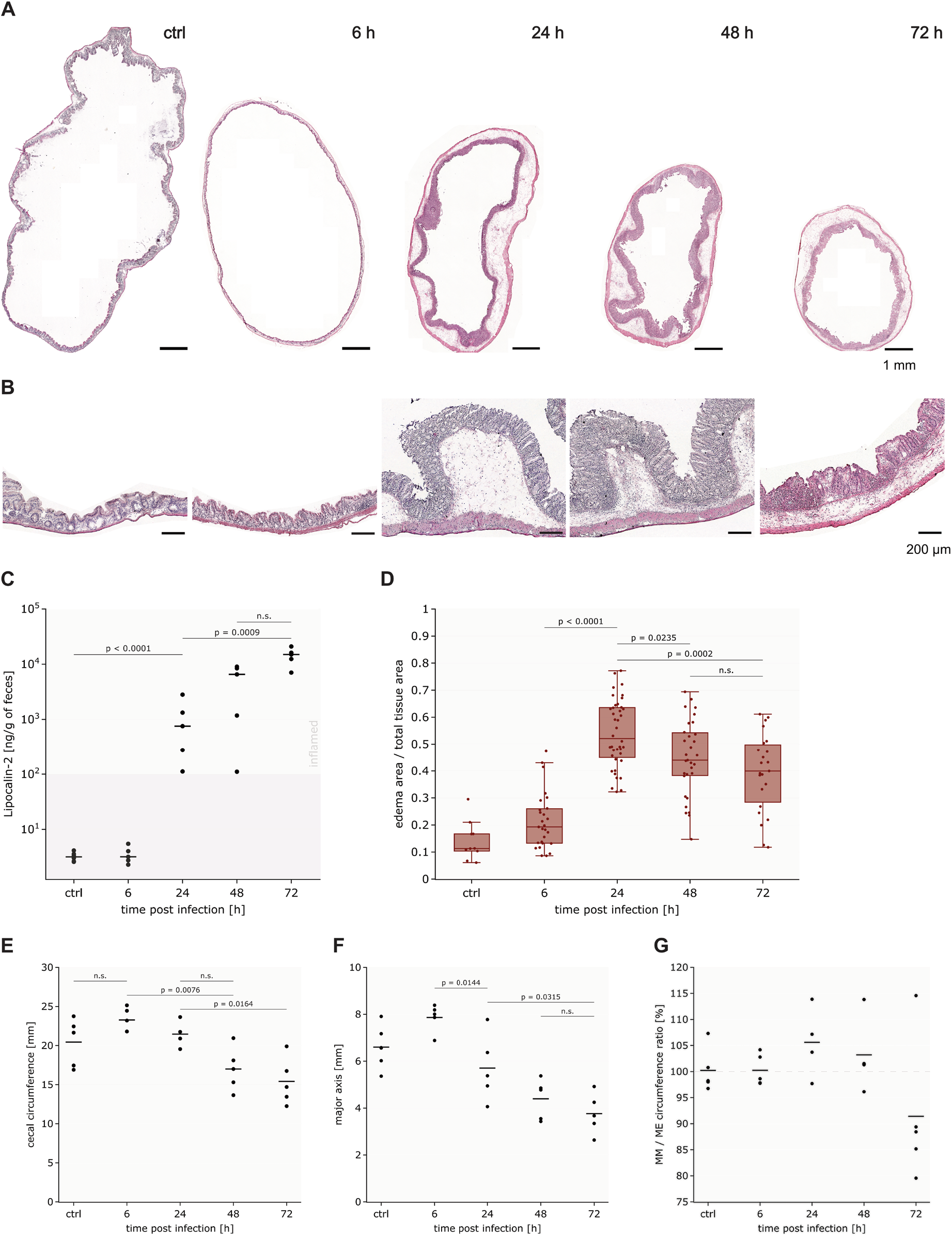
Histopathological analysis of *S.* Tm infected cecum in C57BL/6 mice shows decrease in overall cecum size and cross-sectional luminal diameter with increasing infection time and inflammation. **A:** Representative H&E-stained whole cecum cryosections during the disease time course (control mouse, 6 h p.i., 24 h p.i., 48 h p.i. and 72 h p.i.), reveal morphological changes in all cecal tissue layers. H&E-stained sections for all 25 mice in SFig1. Scale bar: 1 mm. **B:** Higher magnification images of representative parts of H&E-stained cecum sections during the same timeline. After 24 h p.i. the tissues show a clear inflammatory response including submucosal edema formation, polymorphonuclear granulocyte infiltration as well as epithelial integrity loss. Scale bar: 200 µm. **C:** Fecal Lipocalin-2 measurements during the time course of the infection confirm the increase in gastrointestinal inflammation. The dark grey shaded area represents the range of normal values in healthy mice. Marker points represent individual mice (n=5 per group), horizontal lines indicate median of the different groups. Statistical analysis: One-way ANOVA of log-normalized data with Tukey’s multiple comparison test, p-values indicated, n.s.: p ≥ 0.05. **D:** Quantification of the edema fraction of the total tissue cross-sectional area at four different timepoints and in control mice from H&E stained cryosections. Data points represent individual tissue sections analyzed and boxplots extend from the first to the third quartile with median indicated as horizontal line. The whiskers represent the largest/lowest data point of the dataset that falls within 1.5x the interquartile range. **E-G:** Horizontal lines indicate mean value of the different groups. Data points represent averaged values for 4-5 individual mice per group, which were calculated from at least 3 H&E-stained cryosections. Statistical analysis: One-way ANOVA with Tukey’s multiple comparison test, p-values indicated, n.s.: p ≥ 0.05. **E:** Quantification of the outer cecal circumference demonstrates a significant decrease over the course of the inflammation. **F:** Quantification of the major luminal axis represents the strong volume decrease in cecal lumen over the disease course. **G:** Quantification of the ratio between muscularis mucosa (MM) length and muscularis externa (ME) length

### Cecum size and cross-sectional luminal area significantly decreased during inflammation

Edema development was monitored over the course of the disease at 6 h, 24 h, 48 h and 72 h p.i.. All dissected cecum organs were first rinsed with PBS to remove existing intestinal content and parts of the mucus layer in order to avoid detachment of the cryosections from the glass during the staining procedure. Subsequently, the rinsed cecum organs were filled with embedding medium for tissues to preserve the original organ appearance. The intrinsic elasticity and stiffness impeded an overstretching of the organ when manually filling it with embedding medium.

Pathohistological assessment using hematoxylin & eosin (H&E) stains clearly showed the kinetics of tissue remodeling (Fig.1A+B, SFig.1). The circumference of the tissue decreased significantly during disease progression, supporting the visual impression of the cecum size during the dissection of the mice. On the last day of the infection, the circumference of the cecum had decreased by 25% compared to the control group (Fig.1E). This observation can be further validated when looking at the luminal diameter change of the cecum tissue. The major inner axis of a fitted ellipse, representing the diameter of the lumen, decreased from 8 mm in healthy animals to 4 mm in severely inflamed animals (Fig.1F, SFig.2). Furthermore, we analyzed the differences in length between the two muscle layers, muscularis mucosa and muscularis externa from H&E-stained tissue sections as function of time (Fig.1G, SFig.3). In mock infected mice as well as mice at 6 h p.i., the circumference of the inner and outer muscle layer were almost the same. At 24 h p.i., the length of the muscularis externa decreased by 7.5% averaged over multiple sections from the 5 mice (muscularis mucosa decreased by 3.8%). This effect was more pronounced at 48 h p.i. (muscuarlis externa further decreased by 21% and muscularis mucosa by 18%) and the muscularis mucosa layer starts to take on a buckled morphology. This can be clearly seen visually in the H&E-stained cecum cross-section images, showing the mucosa plus muscularis mucosa layer in a buckling formation (Fig.1A+B). The surprising observation was found in mice 72 h p.i., which show a greatly decreased ratio between the inner and outer muscle layer due to a much stronger decrease in muscularis mucosa length relative to the muscularis externa. 4 out of the 5 animals show a much shortened muscularis mucosa length and also morphologically do not show the buckling formation anymore. The one exceptional mouse that still shows a buckled mucosa in the cecal H&E-stained cross-sections and the unaltered muscular ratio, has a lower fecal LCN2 value comparable to the mice on day 2. This indicates that the inflammation in this mouse is a bit delayed and explains the different morphological appearance.

**Figure 2:**
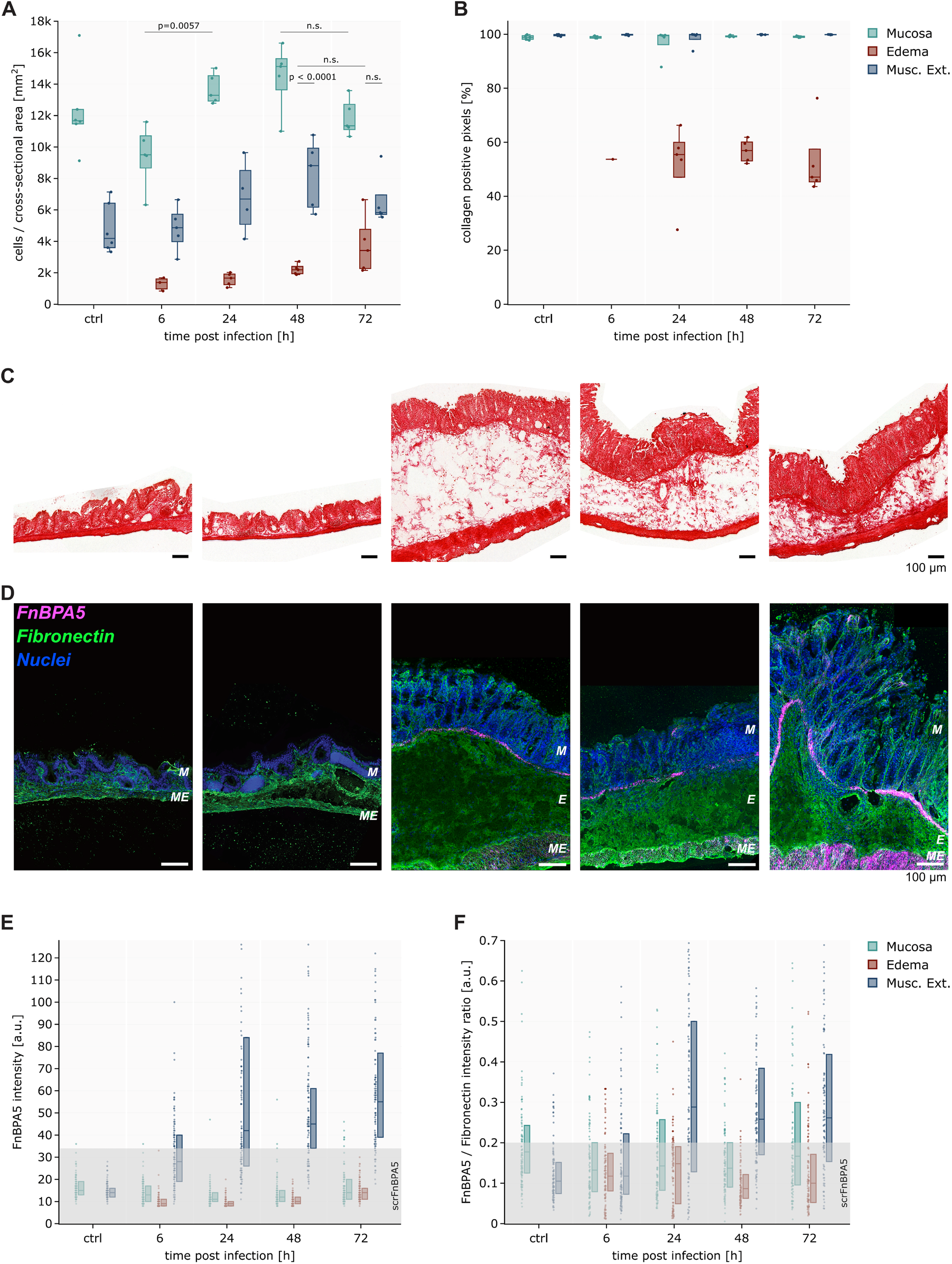
Mechanobiological analysis of the *S.* Tm infected cecum in C57BL/6 mice during disease time course. **A:** Quantification of cell numbers per area in the three different tissue areas (mucosa: green, edema: red, muscularis externa: blue) using H&E-stained cryosections, confirms the low cell density in edema relative to the other tissue layers. Statistical analysis: One-way ANOVA with Tukey’s multiple comparison test, p-values indicated, n.s.: p ≥ 0.05.. **B:** Sparse collagen content in edema areas was quantified by calculating the collagen positive pixel density in the three different tissue areas from picrosirius red stained cryosections. Data points represent averaged values calculated from at least 3 H&E-stained sections per mouse from 5 individual mice per timepoint. Boxplots extend from first quartile to third quartile with median indicated as line. The whiskers represent the largest/lowest data point of the dataset that falls within 1.5x the interquartile range. **C:** Representative picrosirius red stained cecal sections during the disease time course (left to right: control mouse, 6 h, 24 h, 48 h and 72 h p.i.), demonstrate the dispersed collagen signal in edema. Scale bar: 100 µm. **D:** Representative images of immunohistochemistry staining of fibronectin using a polyclonal anti-fibronectin antibody (green), of relaxed fibronectin fibers using Cy5.5-FnBPA5 (magenta) and cell nuclei (blue) during the course of the inflammation (same time points as in C). Additional images from in S.Fig4. Scale bar: 100 µm. **E+F:** Pixel-by-pixel quantification of FnBPA5 intensity and the FnBPA5/Fibronectin intensity ratio, respectively at the same time points. Boxplots extend from first quartile to third quartile with median indicated. A random subset of pixel intensity values (n=100) as individual datapoints presented left to the boxplot. The grey box represents the averaged intensity of the scrFnBPA5 control staining (mean intensity value + 3x standard deviation). ***M***, mucosa; ***E***, edema; ***ME***, muscularis externa.

**Figure 3:**
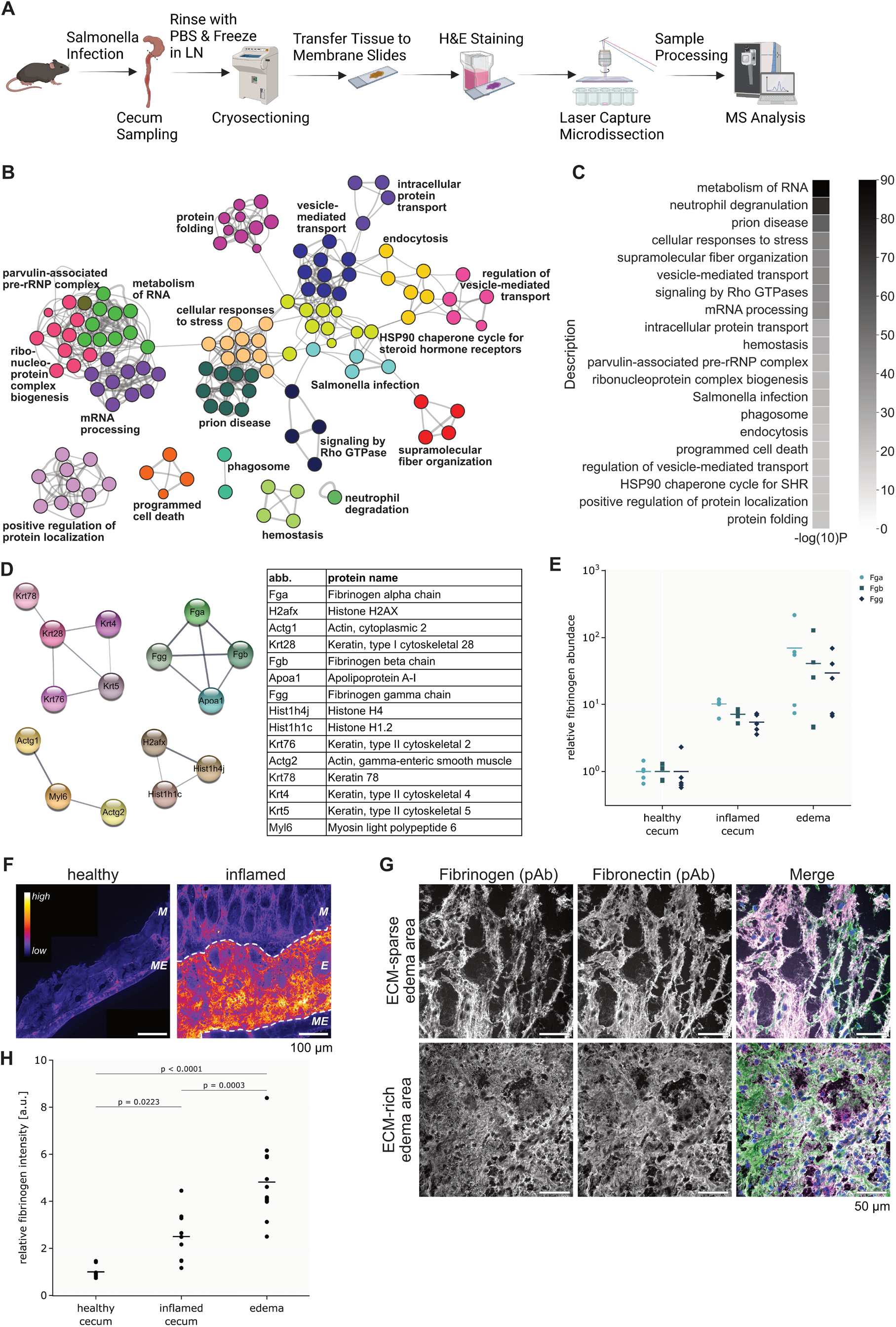
Investigation of the intestinal edema specific proteomic signature in *S*. Tm-infected mice at day 3. **A:** Schematic of workflow for edema-specific proteomics analysis using laser capture microdissection to extract individual tissue areas from H&E-stained cryosections. Illustration generated using BioRender.com. **B:** Overview of all identified Gene Ontology clusters color-coded by similarity. Labels are based on the most representative GO term in the cluster. **C:** Quantification of GO enriched terms of all proteins detected in cecal edema. **D:** Top 15 proteins matched to most abundantly detected peptides represented as network (connecting line thickness scaling with the combined STRING score) as well as table (sorted by detection abundance). **E:** Quantification of fibrinogen abundance from the edema-specific proteomics dataset and comparison with total healthy and inflamed tissue (without edema area) abundance. Fibrinogen alpha (Fga), beta (Fgb) and gamma (Fgg) chain quantified separately (circle, square, diamond markers, respectively). Data points represent individual mice with the mean indicated as horizontal bar. **F:** Representative image of immunohistochemistry staining of fibrinogen in healthy and inflamed (72 h p.i.) cecum sections using a polyclonal anti-fibrinogen antibody. Scale bar: 100 µm. **G:** Representative images of co-staining for fibrinogen (magenta) and fibronectin (green) in a sparse (upper panel) and a less-sparse (lower panel) region of cecal edema. Scale bar: 50 µm. **H:** Quantification of fibrinogen signal intensity in total healthy tissue and relative signal intensity in total inflamed as well as edema areas shows significant increase of fibrinogen intensity in edema tissue. Horizontal lines indicate mean value of the different groups. Data points represent analyzed images from healthy and inflamed (72 h p.i.) mice (9-12 images of at least 3 mice analyzed for the different categories). Statistical analysis: One-way ANOVA with Tukey’s multiple comparison test, p-values indicated, n.s.: p ≥ 0.05.***M***, mucosa; ***E***, edema; ***ME***, muscularis externa.

Specifically focusing on the edema, at 6 h little to no swelling of the submucosa was observable. By the 24 h time point, severe edema formation is clearly visible. Surprisingly, we observed a decrease in edema area as fraction of total tissue area over the remaining course of the disease (48 h and 72 h p.i.). While we observed large edema throughout the 3-day time course, on day 1 of the infection almost 55% of the tissue area was occupied by edema, which decreased to 40% on average on day 3 (Fig.1D).

### Cell numbers and collagen content are significantly reduced in intestinal edema compared to the surrounding tissue layers

Another interesting observation made from the histological assessment was the low number of cells in the submucosal layer in comparison to the mucosa and muscularis externa. In line with expected fluid accumulation, the quantification of cell number showed an almost 10-fold lower cell number per unit area in the edema in comparison to the mucosal layer (Fig.2A). In addition, we observed an increase in cell number in mucosa and muscularis mucosa until day 2 post infection, which is probably caused by infiltration of neutrophils and other proinflammatory immune cells (Maier et al., 2014). On day 3, cell numbers dropped in these two tissue layers likely indicating a terminal loss of tissue integrity. Picrosirius Red staining for collagen proved the high collagen content in mucosa and muscularis externa during the complete time course (Fig.2C). In contrast to the dense collagen network in these tissue layers, the collagen content in the edema is sparse and quantification of positive picrosirius red stained pixels yielded a collagen positive pixel density of only 50% compared to the mucosa and muscularis externa (Fig.2B). The importance of tissue collagen in providing a stable matrix for cells and maintaining tissue integrity is well acknowledged (Diller and Tabor, 2022; Pompili et al., 2021). Thus, the loss of collagen content in the edema tissue might indicate tissue instability and loss of tissue integrity.

### Intestinal edema contain stretched fibronectin fibers while the adjacent smooth muscle layers show major fibronectin fiber relaxation during the infection

Growing evidence supports an active role of extracellular matrix and its remodeling in the process of edema formation (Reed and Rubin, 2010; Wiig et al., 2003). ECM remodeling not only includes stiffer scaffolding proteins such as collagens (Reed et al., 2010), but also elastic fibers, proteoglycans and glycoproteins such as fibronectin, which is known to interconnect parts of the ECM via a plethora of binding partners as well as for its mechanosensitive signaling abilities (Patten and Wang, 2020; Vogel, 2018; Zollinger and Smith, 2017). We used the bacterially derived FnBPA5 peptide to investigate the tensional state of fibronectin fibers in cryosections (Arnoldini et al., 2017). The FnBPA5 peptide binds with nanomolar affinity to the N-terminal fibronectin domains FnI_2-5_ by formation of a network of multivalent backbone hydrogen bonds with the equilibrium structure of those fibronectin modules. In stretched fibronectin fibers, the structural match and thus affinity between FnI_2-5_ and the FnBPA5 peptide is lost as the distance between the fibronectin type I modules is increasing (Hertig et al., 2012). Coupling of the FnBPA5 peptide with a fluorophore thus allows to specifically detect relaxed fibronectin fibers in *ex vivo* animal tissues. This can indicate ECM remodeling processes. Of note, FnBPA5 cannot distinguish between fibronectin fibers that are structurally relaxed due to the presence of other load-bearing ECM fibers, such as collagen fibers, or due to fibronectin fiber cleavage since both processes increase the multivalent binding of FnBPA5 to sequential type I domains. In contrast, polyclonal antibodies bind fibronectin independent of its tensional state. To control for unspecific binding of FnBPA5, a scrambled version of the peptide (scrFnBPA5) was applied.

20 µm thick cryosections of cecum at different time points (6 h, 24 h, 48 h and 72 h p.i.) were thus used to stain for total fibronectin using a polyclonal fibronectin antibody and FnBPA5 coupled to Cyanine5.5. Surprisingly, there was no evidence for FnBPA5 binding in the edema regions at any time point of the disease (Fig.2D). This was unexpected, as inflammation-associated proteases capable of cleaving fibronectin were anticipated to be present in high concentrations in the edema. Quantification of FnBPA5 intensity averaged over the total mucosal areas showed values in the range of the scrFnBPA5 control staining during the complete time course (Fig.2E). This stands in stark contrast to observations in the muscle tissue (muscularis mucosa and muscularis externa). FnBPA5 signal in the muscularis externa significantly increased after 24 h and increased further over the next two days. Quantification of total (stretched plus relaxed) fibronectin intensity showed a generally higher signal in the muscularis externa compared to mucosa and edema. Total fibronectin intensity values remained quite constant during the disease course in all tissue layers (SFig4A). Quantification of the intensity ratio of relaxed fibronectin to total fibronectin to control for variations in the fibronectin intensity levels, corroborates the low levels of FnBPA5 in edema during the disease progression (Fig.2F). The specific absence of fibronectin fiber relaxation in edema contrasts with observations made in tumor models (Fonta et al., 2023, 2020) and suggests the potential role of stretched fibronectin fibers in maintaining tissue integrity in intestinal edema. Thus, we suspected that mechanisms are in place to protect against proteolytic cleavage of fibronectin at these sites. To investigate this further we established a workflow to identify the edema-specific proteome.

### Intestinal edema protein composition revealed by laser capture microdissection coupled to high-sensitivity MS Proteomics

Identifying the composition of intestinal edema is methodologically quite challenging as the contribution of edema to the overall tissue proteome is small. Therefore, laser capture microdissection was employed to extract edema areas from cecum sections of day 3 *S.* Tm infected mice. Edema tissue areas were collected from serial 20 µm thick cryosections to match a total tissue volume of approximately 0.02 mm^3^. This small amount of tissue required an optimized procedure during the sample lysis, protein precipitation and digestion (see materials and methods). The digested samples were loaded onto EvoTip Pure tips and were run on an Orbitrap Exploris mass spectrometer using data-independent acquisition (DIA) mode and a standardized acquisition method (Xuan et al., 2020). The raw files were analyzed using DIA-NN, a software developed to process highly complex DIA data using deep neural networks (Demichev et al., 2020). A graphical schematic of the workflow is presented in Fig.3A.

In total, 3798 unique proteins could be identified from which 1753 were found present in all 5 infected mice investigated. Gene Ontology analysis of these 1753 proteins using Metascape could identify 1534 matched proteins with the Metascape database (Zhou et al., 2019). Reassuringly, the analysis revealed the statistically enriched gene ontology terms “Neutrophil degranulation”, “cellular responses to stress” and “Salmonella infection” amongst other highly significantly enriched pathway terms. To give a general overview of the proteins that were found in the edema tissue, Fig.3B represents a cluster map of the most significantly found GO terms. Interestingly, “supramolecular fiber organization” was the fifth most significantly enriched cluster, highlighting the importance of fiber reorganization in *S.* Tm-induced edemas (Fig.3C).

To deepen our analysis, we then looked into the abundances of individual proteins and protein classes. Importantly, as an edema can only occur in diseased tissue, a relevant comparison to a “healthy control state” is not possible. The submucosal tissue layer in the cecum is barely visible in a healthy, non-infected mouse and thus cannot be analyzed separately from the other tissue layers. Therefore, relative abundances of edema proteins are presented and will be discussed in the scope of already existing knowledge about edema generation, with a major focus on involvement of ECM in this process. The most abundantly detected peptides mapped to the three fibrinogen chains (alpha, beta, gamma) important for fibrin clot formation as well as mediating inflammatory responses, and to the cytoskeletal components actin, keratin type I and II, and myosin (Fig.3D).

The importance of blood coagulation components, with fibrin(ogen) as a key player, is further strengthened by the GO enriched cluster “Hemostasis”. To verify the finding that fibrin(ogen) was found in such high abundances in edema, we performed immunohistochemistry staining on cecal sections from healthy and day 3 *S.* Tm infected mice. A clear increase in total fibrin(ogen) signal in the tissue could be seen in the severely inflamed tissue compared to healthy controls (Fig.3F). The highest fibrino(ogen) intensities were observed in the edema regions, showing 2.5-fold and almost 5-fold higher mean fluorescence intensity between edema regions and the rest of the healthy and inflamed tissue, respectively (Fig.3H). It is important to point out here that this observation cannot be solely explained by posthumous blood clotting, since inflamed and control tissues were handled identically, and we find a significant increase in edemal fibrinogen content from the immunohistochemistry staining as well as the proteomics data between these two conditions. Comparing the abundance of all three fibrinogen chains detected in the proteomics dataset, revealed a 47-fold increase comparing edemal fibrinogen abundances with total healthy tissue (combining mucosal and muscularis externa proteomics data of healthy mice) (Fig.3E). Co-staining with fibronectin revealed a fibrous fibrin-fibronectin matrix with visual similarity to those found in wound healing (Fig.3G).

### Identification of matrisome-specific proteins

Additionally, the proteomics dataset was analyzed with a special focus on extracellular matrix proteins. 125 from the 1753 proteins could be mapped to the matrisome database generated and curated by Naba et al. (Naba et al., 2017). The matrisome can be separated into the core matrisome (45%) as well as matrisome-associated proteins (55%). Fig. 4A shows the distribution of matrisome proteins found in our edema tissues. In the core matrisome we found 63% of ECM glycoproteins, 16% of proteoglycans and 21% collagens, comprised of 12 different collagen family members. Strikingly, the majority of proteins in the matrisome-associated protein group were ECM regulators such as multiple members of the SERPIN family (59%). Serpins, a group of protease inhibitors that inhibit target enzymes by inducing a conformational change, function in hemostasis, since keeping serine proteases’ activities in check is essential for blood coagulation (Silverman et al., 2001), as well as in infection and inflammation (Bao et al., 2018)). ECM-affiliated proteins such as annexins, galectins and cathepsins as well as secreted factors such as the members of the protein S100 family make up 30% and 10% of the matrisome-associated protein class, respectively. Galectins are small carbohydrate-binding proteins with intra- and extracellular functions, where they modify interactions with and within the extracellular matrix, as well as with microbes (Johannes et al., 2018). Cathepsins are the most abundant lysosomal proteases that get released into extracellular space upon tissue damage. The calcium-binding S100 proteins, when released from the cytoplasm upon tissue damage, serve as a danger signal, as they are crucial in the regulation of immune homeostasis, post-traumatic injury, and inflammation (Liu and Stowell, 2023). They can also enhance the adhesion and migration of leukocytes to the site of injury (Xia et al., 2018).

**Figure 4:**
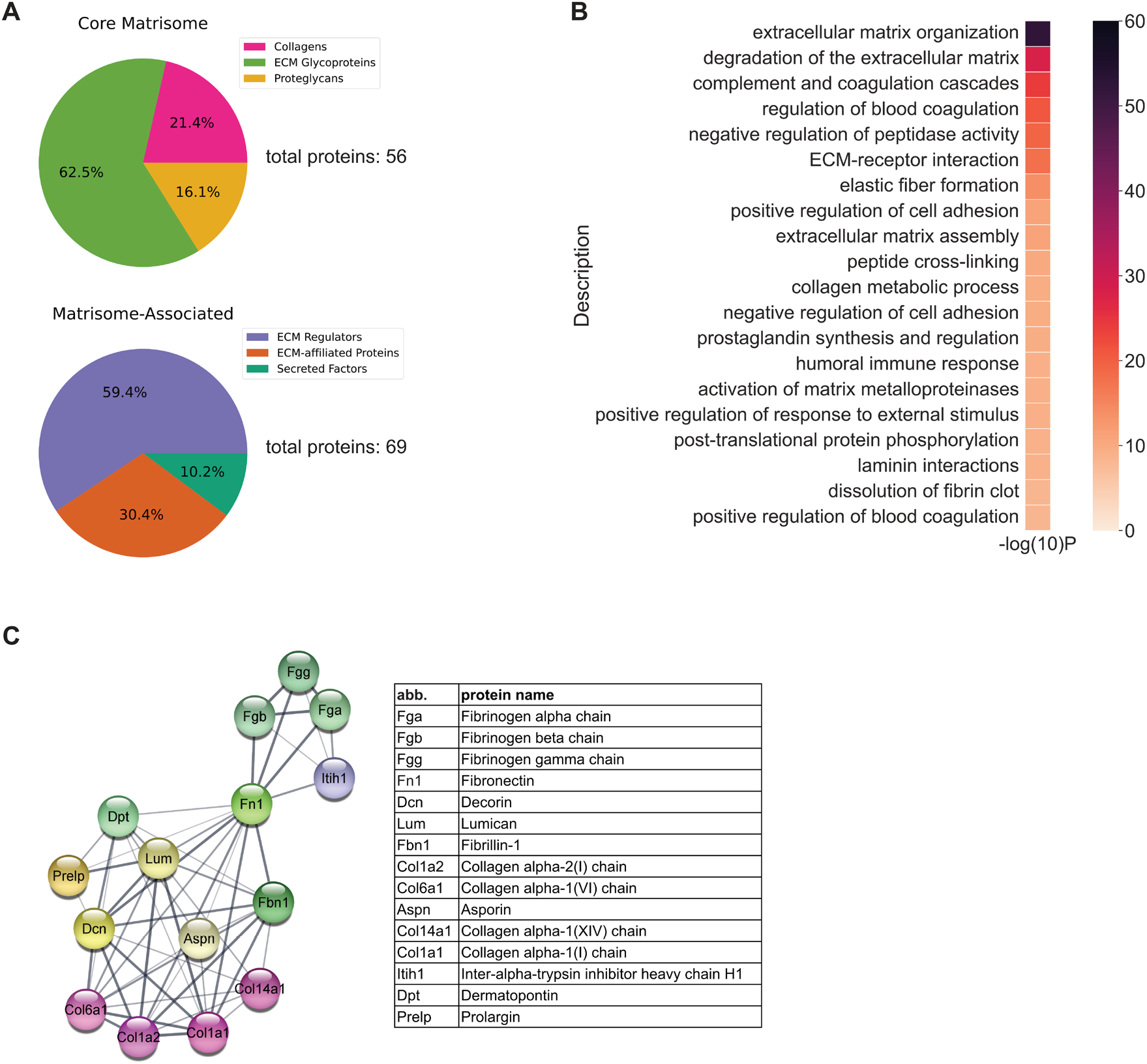
Matrisome-specific protein network detected in *S*. Tm-inflammation induced intestinal edema at day 3 in C57BL/6 mice. **A:** Affiliation of detected edema proteins to the six different matrisome categories. 56 proteins detected are affiliated to the core matrisome category (ECM glycoproteins: 62.5%, collagens: 21.4%, and proteoglycans: 16.1%). In the matrisome-associated category, 69 proteins were detected (ECM regulators: 59.4%, ECM-affiliated proteins: 30.4%, and secreted factors: 10.2%). **B:** Quantification of Gene Ontology enriched terms of matrisome specific proteins detected in cecal edema. **C:** Top 15 matrisome-specific proteins matched to most abundantly detected peptides represented as STRING network as well as table (sorted by detection abundance), highlighting the important role of fibronectin in this network (connecting line thickness scaling with the combined STRING score).

### ECM specific proteins in edema are involved in blood coagulation and negative regulation of peptidase activity

To investigate the functions of these matrisome annotated proteins, an ECM-specific GO enrichment analysis was performed. The top hits were related to extracellular matrix remodeling, regulation of blood coagulation, as well as negative regulation of peptidase activity (Fig.4B). The most abundantly-detected peptides mapped to 15 proteins including fibrinogen, decorin, fibronectin and various collagen chains (I, VI, XIV). Interestingly, the edema specific matrisome network assembles around fibronectin as the core protein, confirming its central role in matrix remodeling and suggesting its important edema-specific role in maintaining tissue integrity (Fig. 4C). The exact influence of the detected matrisome proteins on inflammation is difficult to infer from our dataset, since collagens, decorin and fibronectin all have distinct pro- and anti-inflammatory properties and any interventions will have a narrow window between influencing disease kinetics and causing disastrous loss of tissue integrity, i.e. such interventions go beyond the scope of this study (McQuitty et al., 2020). Nevertheless, the high number of proteins involved in extracellular matrix remodeling is in line with other wound healing and tissue remodeling situations such as skin wound sites (Pfisterer et al., 2021; Potekaev et al., 2021). The regulation of blood coagulation seems to be very important and in fact it would be highly interesting in the future to investigate the role of the coagulation pathway in *S.* Tm infection biology in more detail. Simultaneously, it is very hard to study *in vivo* due to the association of anti-coagulation drugs with gastrointestinal bleeding. In line with our observations, former studies showed strong exacerbation of less excessive tissue inflammation models by anticoagulants (Seltana et al., 2022). Our findings of high levels of fibrin(ogen) together with fibrillar, stretched fibronectin suggest that there is active early wound healing processes ongoing during acute non-typhoidal *Salmonellosis* in the mouse cecal edema.

### Protease and Protease Inhibitor Network in Intestinal Edema

To better understand why mostly stretched fibronectin fibers were detected in edema, we looked further into tissue remodeling processes with a focus on specific proteases and protease inhibitors present in our dataset. For this, we extended the analysis again to the whole dataset (proteins found in edema of all 5 mice, not limited to the matrisome database mapped proteins only). Using the PANTHER database, we allocated protein classes to the protein hits we found (Mi and Thomas, 2009; Thomas et al., 2022). Our data set contains 67 proteins matched to the PANTHER protein class “proteases” as well as 28 proteins allocated to the class “protease inhibitors”. The protease class can be further broken down into 13 cysteine proteases, 13 metalloproteases, 10 serine proteases, 1 aspartic protease as well as 30 unspecified proteases containing mostly proteasome components. The relative abundances with which these proteins were found in the edema of all 5 mice are presented in protein-protein network format in Fig 5. Manual detailed curation of the dataset using the MEROPS database (Rawlings et al., 2014), revealed the presence of several known fibronectin-targeting proteases. Matrix metallopeptidases-3, −8, and −10 were detected in the edema tissues as well as the cysteine protease legumain (Lgmn: Uniprot accession: O89017), which is known to cleave fibronectin between Asn631 + Ala. Interestingly, the reported inhibitor of Lgmn, Cystatin-C (Cst3: Uniprot accession: P21460) was also found with similar abundance. Additionally, we detected the protease inhibitor calpastastin (Uniprot accession: P51125) together with its targets calpain in the protease network. Calpastastin is known for inhibiting calpain 1 (Uniprot accession: O35350) and calpain 2 (Uniprot accession: O08529), which in turn were shown to have fibronectin fragmentation activity (Dourdin et al., 1997; Wan et al., 2016). Overall, this shows multiple protease inhibition possibilities in the *S.* Tm-induced edema to protect fibronectin fibers from being degraded.

**Figure 5:**
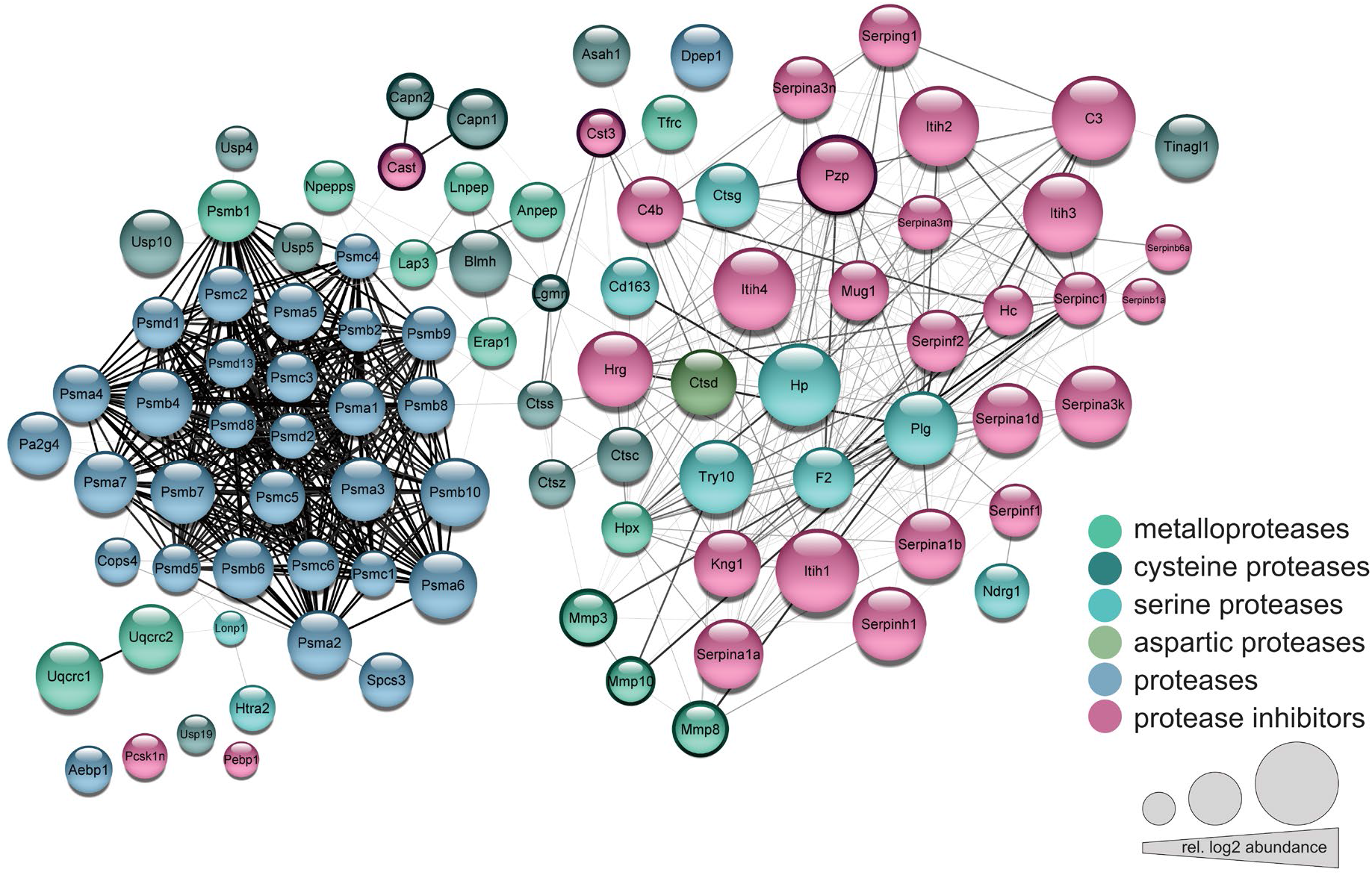
Protease - protease inhibitor network in intestinal edema at day 3 in C57BL/6 mice. Intestinal edema specific protease - protease inhibitor network reveals 67 proteases and 28 protease inhibitors. Multiple proteases with fibronectin cleavage capability were detected together with their protease inhibitors (highlighted with dark border). The circle color represents the class (protease inhibitors, proteases, metalloproteases, cysteine proteases, serine proteases and aspartic proteases) and the circle size the relative log2 abundance with which this protein was detected in the dataset. The grey lines represent protein associations predicted using the STRING database (connecting line thickness scaling with the combined STRING score).

## Discussion

To address the impact of ECM during gastrointestinal inflammation, we characterized cecal tissue alterations during inflammation from a biochemical and mechanobiological perspective. We identified the proteomic signature of intestinal edema in the late stage of the *S.* Tm infection in mice and revealed the presence of tensed fibronectin fibers and of a large protease and protease inhibitor network mostly tracing back to the blood coagulation pathway in intestinal edema.

In general, intestinal tissue is subjected to a plethora of mechanical forces under normal as well as pathological conditions. These physical forces act at all length scales, from the macroscopic to the molecular level, including muscular contractions (Hennig et al., 1999), shear stress, cell-cell interactions, and lymphatic flow (Gayer and Basson, 2009; Malijauskaite et al., 2021; Swartz and Lund, 2012). During tissue inflammation these forces are altered and can induce changes in cellular mechanotransduction processes (Du et al., 2022). Consistent with the previously reported shrinkage of the cecum during *S*. Tm infection (Barthel et al., 2003), we observe a significant reduction of luminal area as well as a change in muscle contraction of the two prominent smooth muscle cell layers, muscularis mucosa and muscularis externa. In the healthy cecum the two muscle layers show a circular structure and are in close proximity to each other. Upon the onset of the inflammation, the muscularis mucosa along with the mucosal layer switch from this circular to a more buckled morphology. The luminal volume loss seems to be steered by both the contractions of the smooth muscle layers and the increasing tissue volume, driven by increased mucosal thickness and edema built up in the submucosa. Reducing the potential volume that luminal *Salmonella* can colonize is an obvious benefit to the host. However, the forces created by massive fluid influx thereby are challenging tissue integrity as the inflammation progresses. The buckling of the inner muscle layer during the first 2 days of bacterial infection maintains anchor points between the inner and outer muscle layers despite edema formation, however, this anchorage is mostly lost on day 3. The circular structure of the inner muscle layer in the presence of edema might thus indicate one step towards full tissue disintegration of the inflamed cecum in *S*. Tm infected C57BL/6 mice, and this infection is typically lethal on day 4/5 in this model (Roy and Malo, 2002).

To ask whether molecular mechanisms might have evolved to reduce the risk of tissue rupture in severe inflammation, we further characterized the molecular composition and ECM properties of *S.* Tm-induced intestinal edema. It contains a loose fibrous network of matrix components, especially a comparatively sparse collagen fiber network, and a limited number of cells. The peptide tension sensor FnBPA5 only binds to low but not to tensed fibronectin fibers, as fiber stretching and partial domain unfolding destroys its multivalent binding motif (Hertig et al., 2012). FnBPA5 staining revealed that fibronectin fiber tension in bacterially infected cecum tissue is dependent on the tissue layer. While fibronectin fiber relaxation is triggered after one day of infection already in the muscle layers (at least in part due to the total decrease of the circumference length of the muscle), the tensed fibronectin fibers in the edema are maintained during the whole course of inflammation. Fibronectin is known for its capability of force-induced extension and contraction, once the tensile forces are released, whereby its extension is associated with structural unfolding which is reversible (Oberhauser et al., 2002; Smith et al., 2007). Fibronectin fibers can expand to more than 6-fold *in vitro* and recover first the original contour length and subsequently their mechanical stability within minutes after release (Klotzsch et al., 2009). Upon stretching, they can thus store elastic energy. On release of applied forces or acting hydrostatic pressure the refolding of the fibronectin fibers will pull the anchor points back into proximity, recovering tissue structure. Due to the low amount of collagen and low number of cells in cecal edema, we hypothesize that fibronectin fibers may actually be the main force-bearing structural component in intestinal edema, perhaps in part counterbalancing the disruptive forces imposed by the build-up of hydrostatic pressure between the inner and outer muscle layer. It is conceivable that fibronectin fibers within a network of fibrin might serve as “bungee” cords keeping the inner and outer muscle layers connected even in locations where massive fluid influx leads to their spatial separation.

Interestingly, the finding of stretched fibronectin fibers in inflamed intestinal edema stands in stark contrast to other pathological tissues transformations, where the tensional state of fibronectin fibers has been mapped by the same tension probe. In tumor tissues, fibronectin fibers are found in a relaxed conformation and are spatially correlated with fibrillar collagen bundles (Arnoldini et al., 2017; Fonta et al., 2023, 2020). Proteolytic degradation, or the increased fibrillar collagen expression in tumor tissues and the reported increase in tissue stiffness could enable fibronectin fiber relaxation, at least in part by collagens that are taking over the role as force bearing structural element (Arnoldini et al., 2017; Kubow et al., 2015). In intestinal edema the collagen fiber network is mostly missing, and fibronectin might be left with the task to carry tissue mechanical stresses. Additionally, studies on virus-infected lymph nodes, showed relaxed fibronectin fibers in the virus-infected swollen lymph nodes as well (Fonta et al., 2020). This is highly interesting since both scenarios take place under inflammatory settings in expanding tissue, but we observe opposing molecular conformations of fibronectin fibers. As recently reported, in inflammation-induced swollen lymph-nodes the conduit system of fibroblastic reticular cells in conjunction with fibrillar collagen is taking over a big part of the tissue tension (Assen et al., 2022; Horsnell et al., 2022; Martinez et al., 2019), which could explain the relaxation of fibronectin fibers in lymph-nodes. In the herein studied inflammation induced intestinal edema, we did not observe evidence for a comparable cellular network to take over tissue tension, but rather found stretched fibronectin fibers.

At the molecular level, it is quite surprising that fibronectin fibers are not cleaved in the inflamed tissue, as the infiltrating tissue fluid is typically rich in proteases. Via proteomics on laser capture dissected mucosal and edema tissues, we identified multiple proteases, which are capable of cleaving fibronectin (MMP-3, MMP-8, Lgmn). Based on our tissue stains, we hypothesize that there are distinct protease inhibitor mechanisms in place protecting fibronectin from degradation, specifically in the intestinal edema. This hypothesis was indeed supported by the detection of multiple members of the SERPIN superfamily, pregnancy zone protein, and cystatin C, which are all broad-spectrum protease inhibitors (Fig.5). Although absolute quantification in untargeted proteomics is not possible, we typically observed higher peptide counts for protease inhibitors than for cysteine-, serine-, and metallo-proteases, consistent with this hypothesis.

Taken together, we would like to propose that fibronectin fibers in the submucosa of the intestine might serve so far unrecognized mechanical functions, i.e. by helping maintaining tissue integrity across regions of edema during intestinal inflammation through mechanical stretching and retraction of fiber tension. As the submucosa swells, the fibronectin fibers interconnecting the two muscle layers are extended as the fluid volume separating the two builds up, thereby keeping the muscle layers connected. Fibronectin’s ability to retract to its original length lets us speculate that these fibers might serve as an energy reservoir for a potential tissue contraction in case of edema regression.

ECM remodeling is an active process during intestinal inflammation, with apparent differences between the different layers of the intestine. By applying laser capture microdissection together with state-of-the-art LC-MS-based proteomics on low amounts of biomass, we could identify candidate proteins involved in the regulation of these processes. By combining these data with fibronectin tensional sensor and microscopy, we could observe the abundance of extended fibronectin and fibrin networks in intestinal edema, and loss of collagen fibers, indicating potential load-bearing by a fibronectin/fibrin-based matrix. We have also identified high levels of both proteases and protease inhibitors in edema fluid, opening the door to generation of protein/gene-based reporters to study edema formation in animal models, and potential pharmacological targeting of edematous intestinal inflammation.

## Materials & Methods

### Ethics statement

All animal experiments were approved by the legal authorities (license ZH120/19) Kantonales Veterinäramt Zürich, Switzerland) and performed according to the legal and ethical requirements. Animals were scored daily for any expected or unexpected adverse events. Humane endpoints were defined in the license. Special focus was put on the 3R principles (Replacement, Reduction and Refinement) for humane animal handling.

### Mice

All experiments were performed with male and female specific-pathogen-free C57BL/6J mice (10-11 weeks old) from an inbred colony at the ETH Phenomics center, and were housed in groups of 2-5 animals in individually ventilated cages in the ETH Phenomics center (EPIC, RCHCI), ETH Zürich. Water and food were given ad libitum during all times. Before the experiments, mice were trained to obtain a fruit-peanut mix (3:2:3 of 20% maltose solution, peanut-oil (sterilized), and fruit puree (fruit puree containing apples (56%), bananas (30% and raspberry (14%), pasteurized)) from the micropipette once per day on two consecutive days. The mice were pre-treated one day before the *S.* Tm infection with 25 mg of streptomycin in 75 µl sterile fruit-peanut solution via voluntary feeding. 24 h later the animals were infected with 5×10^7^ CFU of *S.* Tm (75 µl suspension of sterile fruit-peanut solution) or treated with sterile fruit-peanut solution (control) by voluntary feeding. The *Salmonella* culture was prepared as overnight culture from wild-type *Salmonella enterica* serovar Typhimurium clone SB300, a derivative of strain SL1344 (Barthel et al., 2003), grown for 12 h in Lysogeny Broth (LB) medium containing 50 µg/ml streptomycin at 37 °C. Then the bacterial culture was diluted 1:20 in fresh LB medium without antibiotics and the subculture was grown for 3 h at 37 °C. Before preparing the fruit-peanut suspension, the bacteria were washed twice in cold PBS. At the indicated times p.i. the mice were sacrificed by CO2 asphyxiation followed by blood withdrawal from the heart and tissue samples were dissected for further processing. The mock-infected control group was sacrificed together with the 72 h p.i. group. To avoid cage effects the animals for the individual groups were combined from several cages. During the infection the animals were scored daily.

### Quantification of fecal Lipocalin2

Fecal pellets collected at the indicated time-points were homogenized in PBS by bead-beating at 25 Hz for 2 min. Large particles were sedimented by centrifugation at 300 g for 1 min. The supernatant was analyzed using the mouse Lipocalin-2 enzyme-linked immunosorbent assay (ELISA) Duoset (DY1857-05, R&D Systems) according to the manufacturer’s instruction in serial dilutions.

### Histological procedure

Whole ceca were dissected from the mice and thoroughly rinsed with PBS to remove the content and partially the mucus layer. 3-4 washes with PBS using a rat gavage needle attached to a 20 ml syringe were required to achieve successful washing. In a second step the ceca were filled with cryo embedding medium OCT (#4583, Tissue-Tek® O.C.T. Compound, Sakura) using a rat gavage needle coupled to a 2 ml OCT filled syringe to reach original organ expansion. Depending on the infection state, the volume to fill the cecum varied. Then, ceca were embedded in OCT, snap-frozen in liquid nitrogen and stored at −80°C until further processing. Cryosections for hematoxylin and eosin (H&E) stains and picrosirius red stains were cut at 10 µm thickness. Cryosections for immunohistochemical stains were cut 20 µm thickness (Thermo Scientific Cryostat Microm HM 525). Sections were mounted on glass slides, dried shortly at room temperature, and moved to −20°C for further drying and short term storage. For H&E stains and picrosirius red stains, sections were washed with PBS once before fixed with 4% Formaldehyde solution (#P087.3, ROTI®Histofix, Carl Roth) for 10 min and washed again 2x with PBS and 1x with distilled water. The H&E staining procedure was automated using a histology H&E stainer (COT 20, Medite AG) and samples were embedded using Pertex mounting medium (#41-4014-00, Biosystems Switzerland). For picrosirius red stains, sections were stained with Weigert’s hematoxylin (8 min), washed with tap water, stained with Sirius Red Staining solution (1 hour) and washed twice with 0.2% acetic acid in ethanol (30 sec each). Then sections were dehydrated using 80%, 95% and 100% ethanol solutions (5 min each) and subsequently washed twice in xylol (5 min each) and embed using Eukitt (#03989, Merck).

### Immunohistochemical procedure

Co-stainings for relaxed and total fibronectin fibers as well as for fibrinogen were performed as follows: Tissue sections on glass slides were encircled with a hydrophobic pen (Vector) to decrease staining solution usage. The tissues were then washed once with PBS and blocked with 4% BSA in PBS for 30 min. Cy5.5-FnBPA5 solution was diluted in PBS, added to the sections at a concentration of 5 µg/ml and incubated for 1 hour. For all these steps 100 µl of solution was used per cryosection. The sections were then washed by immersing the whole glass slide into a beaker with PBS (3x 5 min each). In the next step, the tissue sections (20 µm thick) were fixed with 4% formaldehyde solution (#P087.3, ROTI®Histofix, Carl Roth) for 10 min and subsequently washed with PBS 3x for 5 min each. Then sections were blocked for 45 min using a blocking buffer containing 5% goat or donkey serum (depending on the host species of the secondary antibody) (#G9023 and #D9663, Merck) and 0.3M glycine (#56-40-6, Merck) in PBS. Fibronectin was stained using a polyclonal anti-fibronectin antibody (ab23750, Abcam) at a dilution of 1:100 and fibrinogen using the monoclonal anti-fibrinogen antibody (#4440-8004, BioRad) at a dilution of 1:200 incubating over night at 4° C in a humidified chamber. After 3 washes with PBS (5 min each), the secondary antibody goat anti-rabbit IgG Alexa 488 (A11043, Thermo Fisher Scientific) 1:200 (for fibronectin) or donkey anti-sheep IgG Alexa 488 (A-11015, Thermo Fisher Scientific) at 1:200 and donkey anti-rabbit IgG Alexa 647 (A10040, Thermo Fisher Scientific) at 1:200 (for Fibrinogen + Fibronectin) was applied for 1 hour at room temperature. After a quick wash with PBS, co-stain with 4′,6-diamidino-2-phenylindole (DAPI) (D9564, Sigma Aldrich) was performed (10 µg/ml) for 10 min. Last, the sections were washed 3x for 5 min in PBS, dried and mounted using ProLong Gold antifade mounting medium (#P36930, Thermo Fisher Scientific). Mounted and stained sections were allowed to dry at room temperature and stored at 4° C before image acquisition.

### Microscopy

All histological stains were imaged using the automated Slide Scanner Pannoramic 250 (3D Histech). All immunohistochemistry stains were imaged using the Nikon Eclipse Ti2 microscope, equipped with the Yokogawa Confocal Scanner Unit CSU-W1-T2 and operated with the NIS-Elements Software. Images were acquired using a 60x 1.2 CFI Plan Apo VC Water Objective. Z-stack multi-tile images at 3 different locations on the individual cecum section were acquired using a large image scan pipeline and stitched with the NIS-Elements software.

### Quantification of tissue areas

Intestinal tissue layers were identified and segmented using a machine-learning based algorithm in QuPath (Bankhead et al., 2017). The algorithm was trained specifically to detect the three tissue layers muscularis externa, submucosal edema and mucosa. In order to avoid artefacts due to the cryosectioning process, the whole analysis was performed on section from two different parts in the cecum. The first one starting at the dead end of the cecum after trimming through the Peyer’s patch cluster. The second set of section was collected from the middle part of the organ with additional care for the cutting orientation of the organ. In short, three differently oriented sections per mouse were cut (turning the orientation 90° for every section). Cecal circumference, major luminal axis, as well as the ratio between the circumference of the muscularis mucosa and the whole tissue were determined by manual annotations in the CaseViewer Software (Case Viewer 2.4, 3D Histech) designed to support examination of images acquired with the automated Slide Scanner.

### Quantification of cell numbers and collagen content

Images of H&E-stained sections were used for this part of the analysis. The inbuilt QuPath algorithm for cell detection was used on manually selected regions of interest (mucosa, edema and muscularis externa). Picrosirius Red stained sections were analyzed using QuPath. A machine-learning based algorithm was trained to detect collagen positive pixels and then run on the manually selected region of interest representing the three different tissue areas.

### Pixel-by-pixel quantification of FnBPA5 and Fibronectin intensity

FnBPA5 and fibronectin signals were analyzed using a pixel-by-pixel approach. For each mouse at least three different stitched 60x multi-tile images were used at different positions in the cecum and the z-stack maximum intensity projections were used to analyze the images. First, regions of interest in the 3 different tissue areas (mucosa, edema, muscularis externa) were annotated manually in ImageJ (Schneider et al., 2012). Second, all pixel values above the background channel noise were collected for a specific area. Data is represented as boxplots (from all the pixel values for each category) plus additionally randomly selected values (n=100) as marker points indicating the distribution of pixel values left of the boxplots. The grey bar represents the mean intensity (mean of various annotated image regions and mice) plus 3x standard deviation of the control marker scrFnBPA5 and serves as threshold above which FnBPA5 signal is considered specific.

### Mass spectrometry sample preparation and processing

Laser Capture Microdissection was performed using the MMI CellCut Laser Capture Microdissection device (Molecular Machines & Industries). 20 µm cryosections were captured on MMI membrane slides (MMI Prod. No. 50103) and subsequently stained with H&E using the MMI H&E Staining Kit Plus (MMI Prod. No. 70302). The microdissected tissue areas (in total 500.000 µm^2^) were collected using mmi isolation cap tubes (200 µl) and kept at −20° C until further processing. Sample lysis was performed directly on the tube lid. Tubes were opened upside down, fixed in this position and 10 µl RIPA buffer (25 mM Tris•HCl pH 7.6, 150 mM NaCl, 1% NP-40, 1% sodium deoxycholate, 0.1% SDS (Cat# 89900 Thermo Scientific)) were added onto the tissue on the lit. 1 µl of 10x TCEP (Tris(2-carboxyethyl)phosphine hydrochloride (#C4706, Merck) was added to the lid and mixed carefully. 1 µl of CAA (400mM 2-Chloroacetamide (#C0267, Merck) was added, mixed carefully and the tubes were closed. The tissue was incubated for 30 min at room temperature. Liquid was spun down in a tabletop centrifuge and 48 µl ice-cold acetone 100% was added. Samples were incubated at - 20°C overnight. After acetone precipitation, the lids containing isolation caps were exchanged with normal lids to avoid detachment of the silicone inlets during high-speed centrifugation. Samples were centrifuged for 5 min at 21.000 x g. Then the acetone was removed, and the samples were air dried in fume hood. The samples were resuspended in 10 µl 4 M GdCl (Guanidinium Hydrochloride) in 50 mM HEPES (#G3272, Merck) and sonicated in a water bath sonicator for 10 min. For digestion the samples were diluted in 30 µl 50 mM HEPES (pH 8.5), LysC was added in a 1:50 ratio and samples were incubated for 4 hours at 37°C. Afterwards samples were diluted in another 30 µl 50 mM HEPES (pH 8.5), trypsin was added in a 1:10 protease:proteome ratio (w/w) and samples were incubated overnight at 37°C at 350 rpm. Samples were then acidified with trifluoroacetic acid to a final concentration of 1% and pH was verified using pH strips. Subsequently, samples were centrifuged at 21.000 x g for 15 min and transferred onto EvoTip Pure trap columns for desalting and loading the samples onto the LC-MS. The EvoTip Pure tips were used according to the manufacturer’s instructions. In brief, Evotips were rinsed with 20 µl Solvent B (centrifugation at 800 g for 60 s). Then they were soaked in propanol until the Evotips turned pale white and equilibrated by soaking in 20 µl Solvent A (centrifugation at 800 g for 60 s). After loading the samples on the wet Evotips, they were centrifuged again at 800 g for 60 s and washed with 20 µl Solvent A (centrifuged at 800 g for 60 s). Then, 100 µl Solvent A was added and the Evotips were centrifuged again at 800 g for 10 s only.

### Data independent acquisition mass spectrometry analysis

Samples were placed on the EvoSep One liquid chromatography system (EvoSep, Denmark) and analyzed in-line with an Orbitrap Exploris 480 mass spectrometer (Thermo Fisher Scientific) coupled to a FAIMSpro device. Peptides were loaded on a EV1106 C18 column (15cm x 150 µm, 1.9 µm diameter) and separated with the Whisper100 nanoflow and a 20SPD (samples per day) method consisting of a 58 minute gradient. Eluting peptides were injected to the mass spectrometer using a 20 µm fused silica emitter (EV1087), at a static voltage of 2300 V, carrier gas flow of 3.6 L/min and 240 °C ion transfer tube temperature and a positive polarity. A single compensation voltage of −45 V was applied to the FAIMS device during acquisition with a high-resolution MS1 (HRMS1) data-independent acquisition method. MS1 scans were recorded in the orbitrap detector at a 120.000 resolution, with a scan range of 400-1000 m/z, normalized AGC target of 300%, and injection time set to automatic. MS2 scans were recorded over the full m/z range with an isolation window of 8 m/z and 1 m/z window overlap. Peptides were fragmented using HCD (high collision dissociation), with a fixed normalized collision energy of 32%. The orbitrap resolution was set to 60.000 with first mass of 200 m/z. Normalized AGC target was set to 1000%, with maximum injection time set to automatic. MS1 scans were interspersed every 24 scan events (loop count 24), splitting the m/z range in three equal, 200 m/z parts. Raw data were searched with DIA-NN version 1.8, using a library-free (directDIA) approach and MS1 level quantification. The reference proteome was the mouse proteome database obtained from Uniprot (UP000000589 reviewed, accessed 17/06/2022). Precursor FDR was set to 1%, while Met N-terminal excision, Met oxidation and C carbamidomethylation were added as modifications. Match between runs and RT-dependent cross-run normalization were set to True. Trypsin/P was used as the protease, with one allowed missed cleavage, with otherwise default settings. Search results and protein quantification tables were used for further post-processing and analysis.

## Data and Material availability

The mass spectrometry proteomics data have been deposited to the ProteomeXchange Consortium via the PRIDE (Perez-Riverol et al., 2021) partner repository with the dataset identifier PXD043256. Materials available on request to the corresponding author.

## Authors Contributions

The authors contribution to the paper are as follows: study composition and design: R.R., V.V, E.S.; data collection: R.R., K.K.; contribution of analysis tools and expertise for the proteomics part: U.K., K.K.; draft manuscript preparation: R.R., E.S.; All authors provided critical feedback to the manuscript.

## Acknowledgments

We thank Maike Wennekers Nielsen and Marie Vestergaard Lukassen at DTU for their technical support as well as Till Wüstemann for technical support at ETH Zurich. We thank Gianna La Regina and Stefanie Oswald for their great help during their semester projects. We also thank Valdemaras Petrosius and Erwin Schoof for mass spectrometry guidance and optimization of the in-house HRMS1 method. We gratefully acknowledge the Scientific Center for Optical and Electron Microscopy (ScopeM) of ETH Zurich, the ETH Phenomics center, and the Proteomics Core Facility at DTU for their support.

This work was funded by NCCR Microbiomes, a research consortium financed by the Swiss National Science Foundation (E.S.); Swiss National Science Foundation (40B2-0_180953, 310030_185128) (E.S.), European Research Council Consolidator Grant (NUMBER 865730-SNUGly) (E.S.), Botnar Research Centre for Child Health Multi-Invesitigator Project 2020 (BRCCH_MIP: Microbiota Engineering for Child Health) (E.S., V.V., R.R.). R.R. additionally was supported by the EMBO Scientific Exchange Grant (#9683). U.K. and K.K. are supported by a Novo Nordisk Foundation Young Investigator Award (NNF16OC0020670) and acknowledge funding from PRO-MS: Danish National Mass Spectrometry Platform for Functional Proteomics (grant no. 5072-00007B). The funders had no role in study design, data collection and analysis, decision to publish, or preparation of the manuscript.

## Notes

### Competing Interest Statement

The authors have declared no competing interest.

