## Supplementary Figures for "*Salmonella*-driven Intestinal Edema in Mice is Characterized by Tensed Fibronectin Fibers"

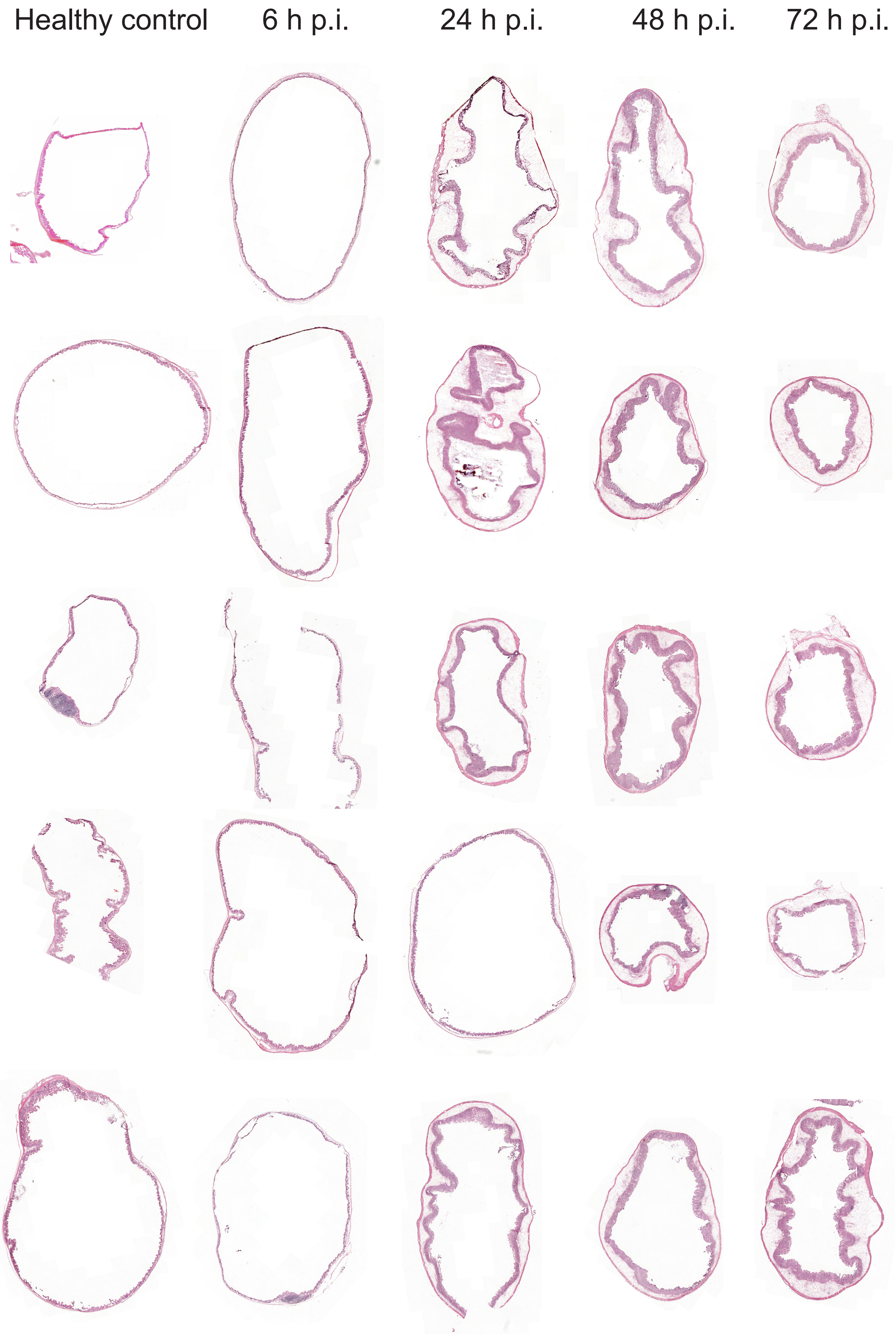


**SFig.1: Overview of H&E-stained cryosections from cecal cross-sections from all 25 mice.** Representative images chosen from several different stained cross-sections. Magnification the same for all imges.

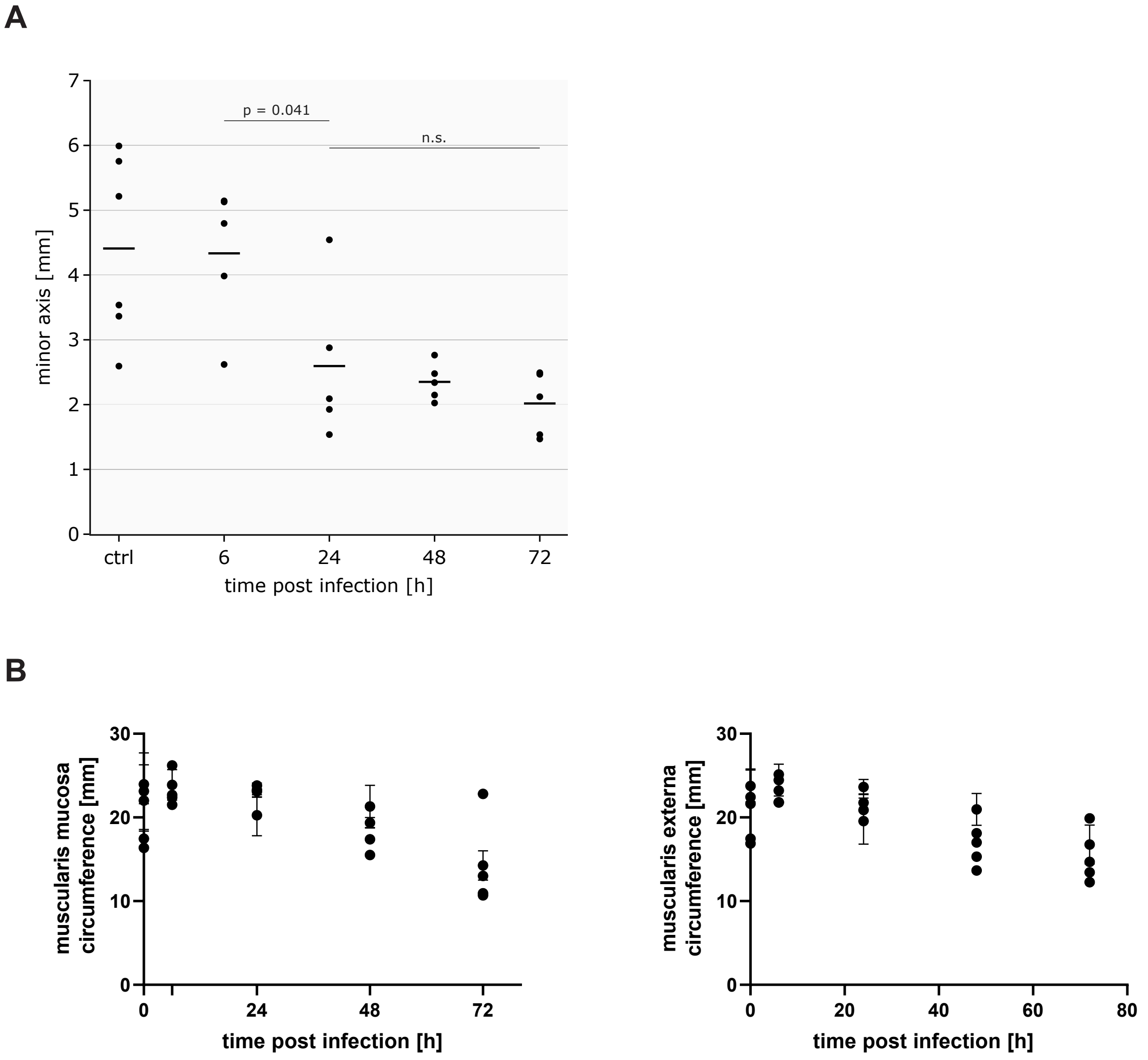


**SFig.2: Quantification of H&E-stained cross-sections. A:** Length of minor axis fitted to the infected cecum lumen. **B:** Muscularis mucosa and muscularis externa circumference quantification individually.

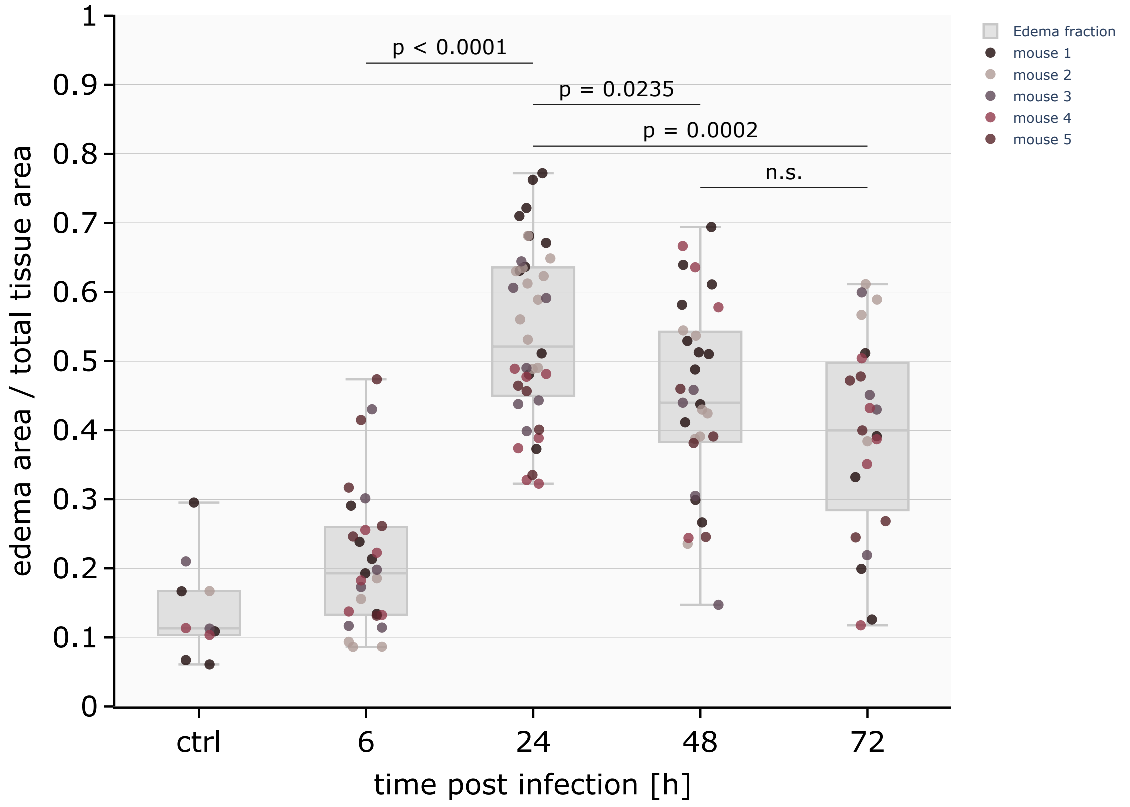


**SFig.3: Quantification of the cross-sectional edema area as fraction of overall tissue area.** The individual mice are shown in different marker colors.

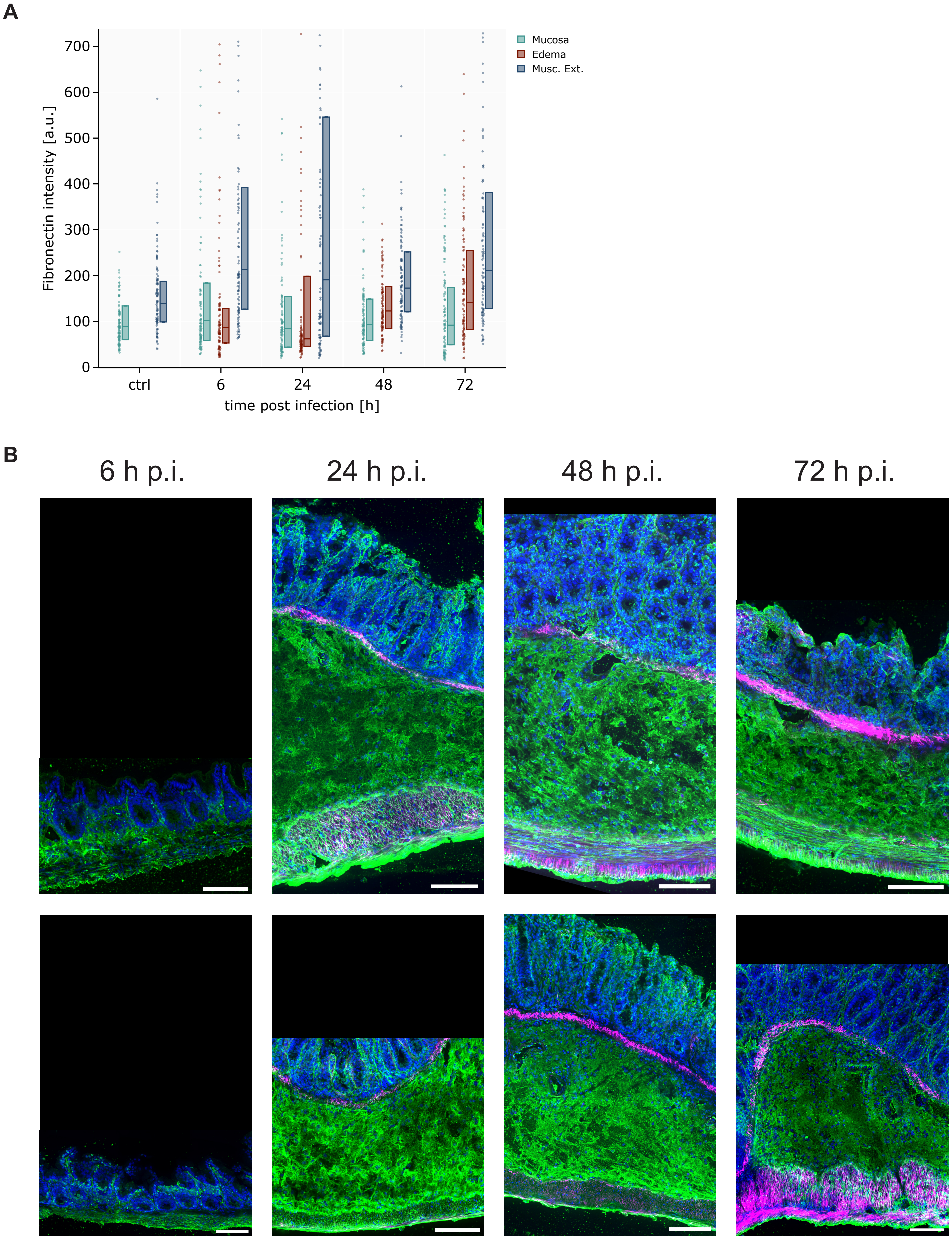


**SFig.4: Fibronectin intensity quantification and additional images showing the FnBPA5 distribution in inflamed mouse cecum. A:** Pixel-by-pixel quantification of the fibronectin intensity over the time course. **B:** Additional images presenting only stretched fibronectin in edema and relaxed fibronectin fibers in the muscle layers and the mucosa. Scale bar: 100 µm.
